# A Theory of Thalamocortical Loops in Evidence Accumulation and Decision-Making

**DOI:** 10.64898/2025.12.23.696263

**Authors:** Michael J. Berry

**Affiliations:** Princeton University

## Abstract

A prominent property of the neocortex is that it forms reciprocal excitatory loops with the thalamus. Here, we propose a function for the loops between layer 5 pyramidal tract neurons in the cortex and matrix-type relay cells in the thalamus, which is based on known anatomy and biophysics. This pathway forms an amplifier circuit with a gain less than unity in the resting condition. However, descending cortical input to apical tufts of L5 neurons can raise the gain of these neurons, boosting the thalamocortical loop into either an integrating or an amplifying regime. Multiple thalamocortical loops compete via lateral inhibition in the thalamus, resulting in a winner-take-all global dynamic among the motor commands encoded by each loop. When L5 neurons are driven by sensory inputs, the integration regime closely resembles the drift-diffusion model of decision-making. We explore basic relationships among parameters of this model and compare broadly against neurophysiological data.

## Introduction

Among the seven major brain divisions, the most mysterious is arguably the thalamus. This mystery stems in part from the fact that the thalamus forms strong recurrent excitatory loops with the neocortex, making analysis methods that have been successful for feedforward pathways less useful. In addition, the thalamus participates essentially in the other two major recurrent loops with the neocortex – namely, the loop from cortex to basal ganglia to thalamus and back to cortex, as well as the loop from cortex to cerebellum to thalamus and back to cortex. Another complexity in understanding the thalamus is the fact that there are important differences in its organization among different thalamic nuclei as well as for cell types within the same nucleus. These anatomical differences are likely to result in substantial differences in the overall function of different thalamic nuclei. In this paper, we propose a computational model that describes the interaction of the neocortex with a major subset of the thalamus, and we analyze its properties.

### Divisions of the Thalamus

Overall, the output of thalamus sent to the neocortex consists of excitatory relay cells that can be divided into three major types – core, matrix, and intralaminar (Clasca et al., 2012; Jones, 2001). Core-type relay cells are parvalbumin positive and project strongly and focally into layers 4 and 3 of the neocortex (Jones, 1998). Core cells are generally considered a relay of sensory information; major examples include the magnocellular and parvocellular relay cells in the lateral geniculate nucleus (LGN) that convey visual information to the neocortex (Jones, 2001). Matrix-type relay cells are calbindin positive and project strongly into layer 1 of the neocortex, as well as several other layers. This is the cell type that we consider in this paper. Intralaminar-type relay cells, also calbindin positive, send a major collateral to the striatum and project to layer 5 and 6 of the neocortex (Clasca et al., 2012). Among other roles, intralaminar cells provide the link between brainstem motor circuits and the basal ganglia for the extrapyramidal motor system (McHaffie et al., 2005); we will not consider them here. Clasca has made a further distinction among matrix cells into one subtype that is focal versus another that is multi-areal (Clasca et al., 2012). Focal matrix cells provide a spatially local to layer 3 of a single cortical area as well as the diffuse input to layer 1 that is typical of matrix-type relay cells. Multi-areal matrix cells provide input to layer 5 as well as layer 1 and have several subcortical branches that connect to multiple cortical areas. The properties of our model most closely match the multi-areal matrix cells (Figure 1A).

**Figure 1.**
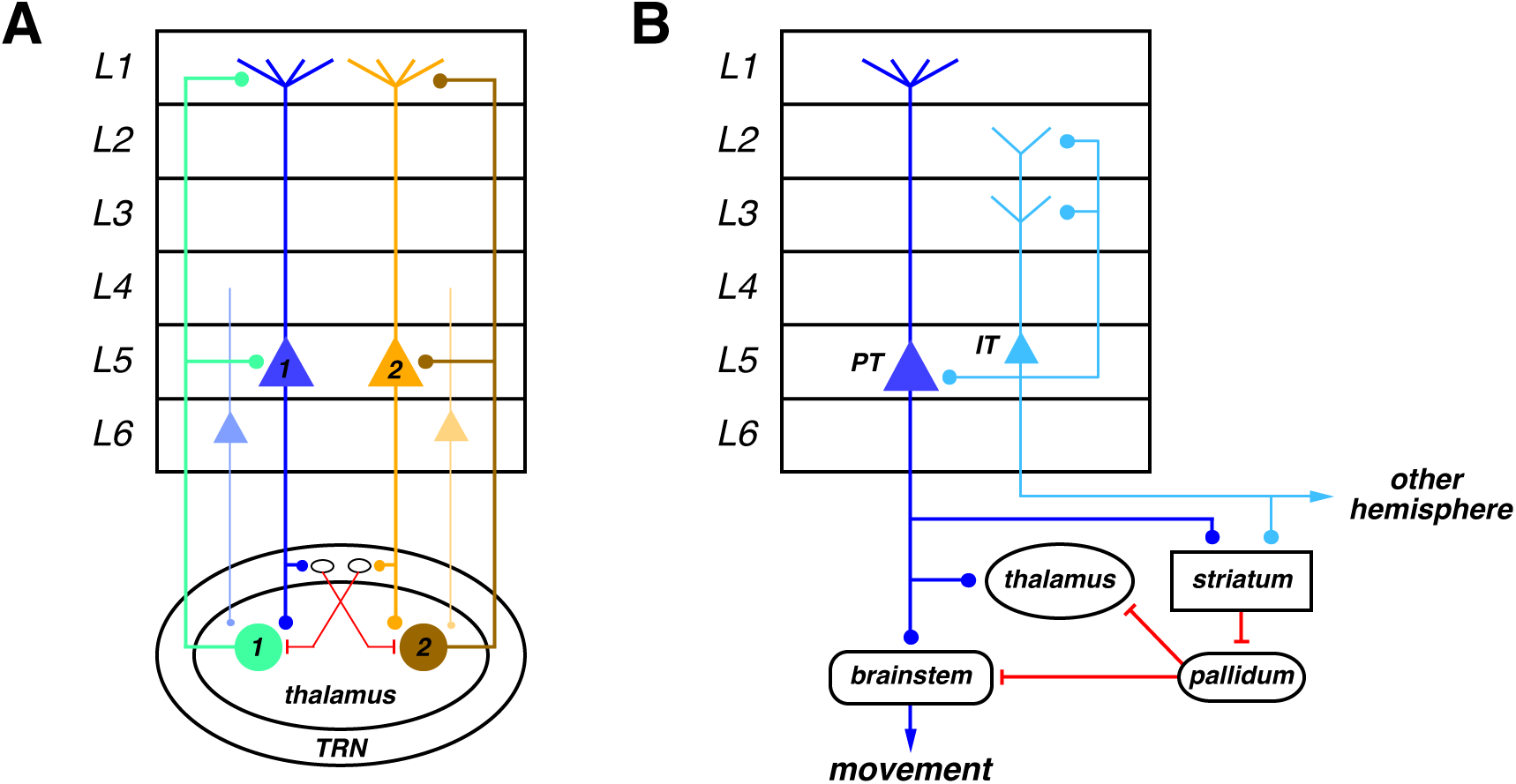
Anatomy of Thalamocortical Loops. **A.** Two populations of layer 5 PT neurons (1: blue, 2: orange) send excitatory driver-type synapses to two populations of multi-areal matrix-type thalamic relay cells (1: green, 2: brown). The thalamic relay cells feed back to the neocortex in layers 1 and 5. The layer 5 PT neurons also send collaterals into the thalamic reticular nucleus (TRN), synapsing onto neurons (black ovals) than make inhibitory contacts onto thalamic relay cells (red). Two populations of layer 6 neurons (lighter shades of blue and orange) send modulatory-type synapses into the thalamus. **B.** Layer 5 PT neurons (blue) project excitation down the pyramidal tract to the brainstem and spinal cord to generate movements. These same neurons send excitatory collaterals into the thalamus and striatum. The striatum transiently inhibits (red) the tonically active pallidum, which pauses its active inhibition to release a movement. Layer 5 IT neurons (light blue) feed excitation forward onto L5 PT neurons, feed back to layers 2 and 3, project down to the striatum, and project through the corpus callosum to the other cerebral hemisphere.

The thalamus receives two different kinds of excitatory afferent fibers: 1) driver-type inputs that form strong ionotropic synapses made onto the soma and proximal dendrites of relay cells and 2) modulator-type inputs that form weak synapses onto the distal dendrites with a large metabotropic glutamate conductance (Jones, 2002; Sherman, 2016). Modulatory inputs come from layer 6 of the neocortex. Driver inputs can be divided into two categories that define two kinds of thalamic nuclei. First-order nuclei receive their driver-type inputs from subcortical sources, while higher-order nuclei receive driver inputs from the neocortex (Sherman, 2017). The model described here pertains to higher-order thalamic nuclei (see Figure 1A).

Because there are no recurrent excitatory connections within the thalamus (Jones, 1981), the circuit performs primarily feedforward computations and thus has often been thought of as a relay. However, as we will see below, the excitatory feedback to the neocortex can give rise to much richer computations and dynamics. In addition, there are two sources of inhibition onto thalamic relay cells: 1) local inhibitory interneurons; 2) the thalamic reticular nucleus (TRN). Both of these sources create lateral inhibition within the thalamus (Hirsch et al., 2015; Wimmer et al., 2015). The excitatory inputs from L5B neurons will excite a specific subset of relay cells monosynaptically while simultaneously inhibiting the other relay cells disynaptically via local inhibitory interneurons. The nucleus reticularis is driven by excitatory inputs from L6 of neocortex as well as axon collaterals of thalamic relay cells (Jones, 2002; Sherman, 2005). This pathway is functionally similar to the action of local inhibitory interneurons but could have greater spatial extent. Recent studies have reduced the estimates of the number of local inhibitory interneurons, suggesting that the dominant source of inhibition is probably via the TRN (Evangelio et al., 2018). Together, these sources of lateral inhibition constitute the primary local computation within the thalamus (see Figure 1A).

### Neocortical Motor Control

All areas of the neocortex possess neurons in layer 5 that project down the pyramidal tract to the brainstem and spinal cord (Prasad et al., 2020; Shepherd, 2013; Swanson, 2000). These neurons are variously referred to as “pyramidal tract” or “PT” due to their projection, “tufted” pyramidal cells due to their morphology, or “layer 5B” due to their laminar position (Thomson & Lamy, 2007) (Ramaswamy & Markram, 2015). These terms all refer to substantially the same set of neurons, which we will subsequently refer to as “L5 PT”. Other studies describe subtypes within this population, but we will not distinguish subtypes here (see Figure 1B). Because the axons of L5 PT neurons project down the pyramidal tract, we interpret their activity as encoding motor commands (Deschenes et al., 1994; Guillery & Sherman, 2002; Sherman, 2017). This interpretation implies that all areas of the neocortex can generate motor commands. At the same time, different areas of neocortex have different relative fractions of L5 PT neurons as well as different specific projection targets. For instance, the primary motor cortex, M1, has the thickest layer 5 and L5 PT neurons with some of the largest somas in the brain (Nolan et al., 2024; Wagstyl et al., 2020), projecting widely to the brainstem and spinal cord. In contrast, the primary visual cortex, V1, has a thinner layer 5, from which L5 PT neurons project primarily to the superior colliculus (Hubener et al., 1990). In fact, microstimulation of the deep layers of V1 triggers saccadic eye movements (Tehovnik et al., 2002).

The axons of L5 PT neurons branch and project to several other targets in the brain: 1) the thalamus; 2) the striatum; 3) pontine nuclei which feed mossy fibers into the cerebellum (see Figure 1B; (Prasad et al., 2020; Ramaswamy & Markram, 2015; Thomson & Lamy, 2007). As described above, the inputs to the thalamus are driver-type and feed onto relay cells in higher-order thalamic nuclei that in turn feed excitation back to the neocortex. Thus, this pathway forms a reciprocal excitatory loop. The inputs to the striatum drive medium spiny neurons (MSNs), which inhibit ongoing activity in the globus pallidus internal segment and substantia nigra pars reticulata (which we together refer to as the “pallidum”). The pallidum projects both to brainstem motor nuclei and higher-order thalamus, providing sustained inhibition to those targets. When medium spiny neurons fire, they pause the firing of pallidal neurons, disinhibiting thalamic relay cells, which in turn provides excitation back to the neocortex. Thus, the projection of L5 PT neurons to the striatum forms another reciprocal excitatory loop. The projection of L5 PT neurons to the cerebellum forms a third reciprocal excitatory loop via feedback from deep cerebellar nuclei to the thalamus and then to the neocortex. However, we will only consider here the role of the primary thalamocortical feedback loop.

The fact that every area of the neocortex can simultaneously generate its own motor commands creates a substantial problem for the brain – namely that these different motor commands need to be coordinated to create a single movement rather than an inconsistent combination of muscle contractions from multiple movements. Furthermore, this coordination needs to be widely consistent across the entire neocortex. We propose here that one of primary functions of higher-order thalamus is to achieve this global coordination among neocortical motor commands.

More specifically, we assume that each movement is encoded by the activity of a specific set of L5 PT neurons, which we refer to as a “pool”. These neurons drive a corresponding set of thalamic relay cells, which feed back to the neocortex, constituting a reciprocal excitatory loop that we refer to as a “TC loop”. We will show below that under specific conditions for the cortical input to a given pool of L5 PT neurons, a TC loop can generate self-sustaining activity that either ramps or increases superlinearly in time. Each TC loop then competes with other TC loops via lateral inhibition in the thalamus. The TC loop with the strongest activity will suppress other TC loops, disinhibiting itself, until its activity reaches the threshold to initiate the movement that it encodes (Figure 1). The dynamics of these interacting TC loops then constitutes a system of global movement coordination via inhibitory competition in the thalamus.

Another key element of neocortical motor control is the fact that the pallidum tonically inhibits brainstem motor nuclei (see Figure 1B; (Grillner et al., 2005)). As a result of this inhibition, L5 PT neurons cannot trigger a voluntary movement merely by sending excitatory signals to the brainstem. In addition, L5 PT neurons must also provide strong enough inputs to the striatum to pause the pallidum and release this “inhibitory brake”. The most vivid demonstration of this requirement comes from late-stage Parkinson’s patients, who are effectively paralyzed because they are unable to overcome the tonic inhibition produced by the pallidum.

In this context, the intrinsic electrical excitability of medium spiny neurons is highly significant. MSN activity in the striatum is observed to be sparse (Barbera et al., 2016; Markowitz et al., 2018). MSNs also express a strong inward-rectifier K^+^ conductance. This ion channel opens when the cell membrane hyperpolarizes, increasing the K^+^ conductance, which acts to further hyperpolarize the membrane. This conductance therefore creates a positive feedback dynamic that drives the membrane voltage towards the K^+^ reversal potential, which is experimentally observed as a “down state” (Wilson & Kawaguchi, 1996). The ultimate result is that the threshold for excitatory inputs to overcome the downstate and elicit action potentials is different and can be considerably higher than the threshold to fire other neurons in this pathway (Steephen & Manchanda, 2009). We model this threshold with the parameter, Θ (Figure 2).

**Figure 2.**
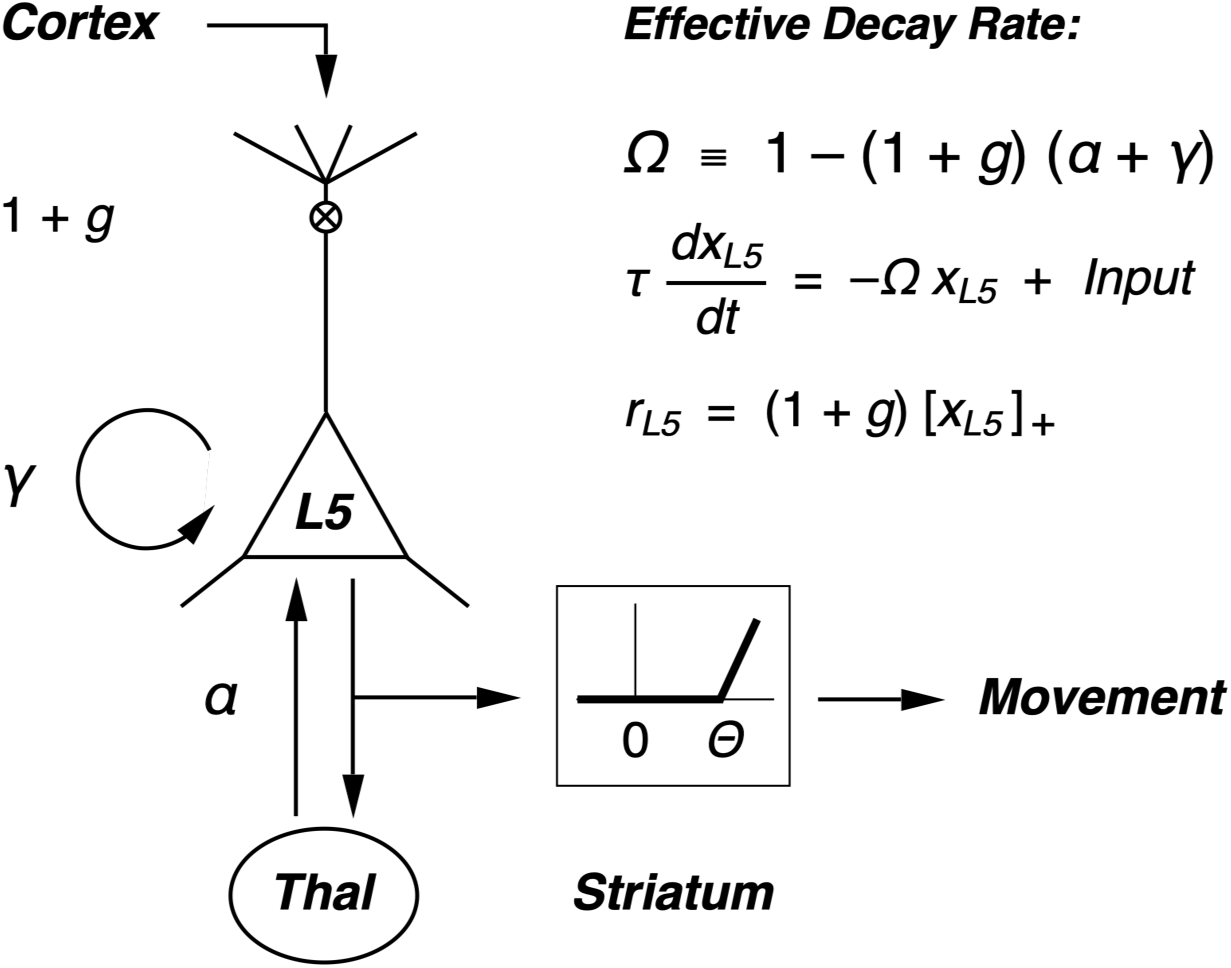
Model of the thalamocortical loop. ***Left***: The thalamocortical (TC) loop consists of a pool of L5 PT cortical neurons that activate thalamic relay neurons. The thalamus feeds back to the same L5 PT neurons with coupling strength, α. L5 PT neurons excite themselves with coupling strength, γ. Finally, descending cortical inputs determine the gain, 1+*g*, of the L5 PT neurons. ***Right***: The dynamics of the TC loop are determined by an effective decay rate, *Ω*, that depends on the parameters, α, γ, and *g*.

The elevated threshold to fire MSNs creates the opportunity for the brain to generate activity in L5 PT neurons that is sufficient to drive most downstream targets, but which is below the threshold to activate the striatum and thereby trigger a movement. We interpret this regime as “motor planning”.

### Structure of the Computational Model

For each thalamocortical loop, there is a population of tufted pyramidal cells in layer 5 that encodes the activation of one specific movement. To the extent that a given movement requires a sequence of motor commands, this pool does not encode for that entire trajectory. Instead, it should be thought of as the initiator of the movement sequence (see Discussion). We model this pool with a single rate variable, *r_i_*(*t*), which should be thought of as the summed firing of all the neurons in pool *i*. It is worth noting that this assumption is not made merely for simplicity. Instead, if there is sufficient redundancy among L5 PT neurons (Narayanan et al., 2005), then the population will have well-defined clusters of activity and activity patterns within the same cluster vary primarily in their total spike count (Berry & Tkacik, 2020).

These L5 PT neurons have two primary inputs: i) basal dendrites in layer 5; ii) the apical tuft in layer 1. For basal dendrites, the input is integrated with a time constant and some decay to produce the “voltage” variable, *x_i_*(*t*):

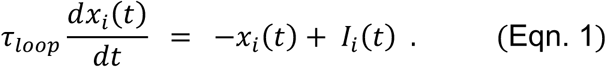

Here, *I_i_*(*t*) represents the total synaptic input to the cortical neurons in pool *i* (see below for details). This input will consist of three main sources: i) feedback from the thalamus; ii) local recurrent input; iii) sensory information (see Eqn. 4 below). This voltage is passed through a nonlinear function to determine the firing rate of the pool, *r_i_*(*t*):

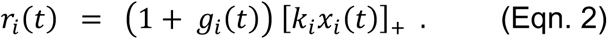

Here, *k_i_* represents a “slope” for the transformation of voltage into firing rate, which has a constant value; *g_i_*(*t*) is the time-varying gain (see below). Without loss of generality, we choose *k_i_* = 1. Finally, the operation, […]_+_, stands for rectification, meaning that negative values are truncated to zero and positive values are unaffected.

For the apical tuft, there is direct evidence that activity in this region effectively sets a “gain” for the neuron (Larkum et al., 2004; Larkum et al., 1999). Thus, inputs into layer 1 will have an instantaneous value that sets the time-varying gain, *g_i_*(*t*). This gain then multiplies the firing rate of the pool, rather than adding to it. Notice also that *g*(*t*) is defined so that zero gain corresponds to multiplication by 1. In general, inputs to layer 1 come from hierarchically higher stages of the neocortex, as well as from the higher-order thalamic nuclei.

The L5 PT neurons in each pool provide input to the thalamus. These are “driver” inputs that make strong synapses onto thalamic relay cells. Furthermore, the thalamus has no local excitatory recurrent inputs. As a result, the firing rate of the downstream pool of thalamic relay neurons, *Th_i_*(*t*), is set every time step by this driving input. However, the thalamus does have lateral inhibition between different pools of relay cells. This lateral inhibition has strength, *w_cross_*, which is a constant. Putting this together, we get:

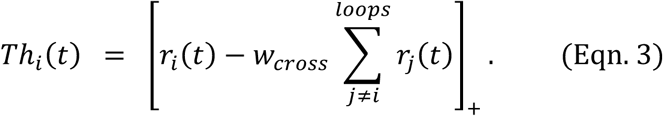

The activity of thalamic relay cells feeds back to layer 5 of the same cortical area that sent it input. These thalamocortical synapses initially target a random set of neurons in layer 5 – not necessarily the specific neurons that provided them input. However, the synapses have Hebbian plasticity, which, over time, will cause many of the synapses onto the same L5 PT neurons to strengthen (Guo et al., 2020; Guo et al., 2018). In the model, we assume that this process has previously occurred. We capture this degree of specific feedback by a single constant, α.

Finally, neurons in layer 5 form an extensive recurrent network, in which local synapses are also shaped by Hebbian plasticity. As a result, neurons within a given pool will form synapses onto other neurons in the same pool. At the level of a single activity variable, we model this as feedback that is proportional to the activity in the pool, γ *r*(*t*), where γ is a constant that characterizes the degree of feedback in the local layer 5 network. This feedback adds persistence to activity in the pool.

As mentioned before, the input to each pool of L5 PT neurons has three main terms, thalamic feedback, local recurrent input, sensory information (Figure 2):

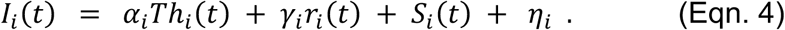

Here, *S_i_*(*t*) is the sensory input to pool *i*. Initially, this input will be zero. The variable η is a spontaneous input that is constant in time; this is included because cortical neurons always exhibit a certain level of random, spontaneous firing. We also note that in principle η could have a component that is due to other inputs from elsewhere in the brain that serve other computational purposes. Notice that the feedback parameter, α, sets how strongly thalamic activity couples back into the pool of L5 PT neurons and that the feedback parameter, γ, sets how strongly the pool of L5 PT neurons activates itself.

The last element that we need in this model stems from the idea a given thalamocortical loop is typically not very active. Such a loop will only be “activated” when its corresponding movement is one that the brain might want to carry out, in which case, the possible movement contributes to motor planning. More specifically, this process is implemented by feedback from higher cortical areas to layer 1 that activates the loop. Recall, that input to layer 1 sets the gain of neurons. Therefore, the gain variable will be set by a control function, *C_i_*(*t*), that represents this cortical feedback:

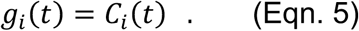

## Results

We begin by studying several simple limits of this model, where we can obtain useful analytical results. First, we consider a single TC loop, next we consider two competing loops, then we consider the case of two competing TC loops driven by sensory stimuli, and finally we consider the effects of additional synaptic pathways.

***1) Single loop with linear dynamics***. Here, we assume that thalamic activity is above the “baseline” level of inhibition in the thalamus, so there is no truncation in firing. We also start by considering only a single loop. Thus,

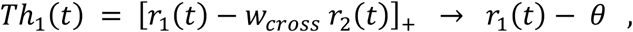

where *θ* captures the effects of other activity in the thalamus and is assumed to be a constant, for simplicity. The feedback to the cortex is then:

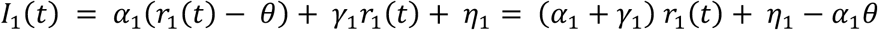

Then the voltage in the cortical pool is expressed as:

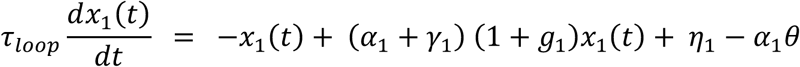

Rearranging terms and dropping the subscripts, we get:

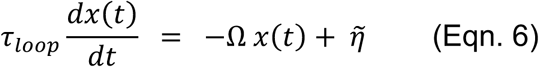

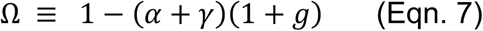

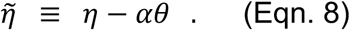

This equation has simple dynamics. In the case of Ω > 0, activity rises exponentially towards a steady-state level, as illustrated in Figure 3A. This happens when the intrinsic decay is larger than the feedback excitation. The steady-state level and dynamics to approach steady-state are:

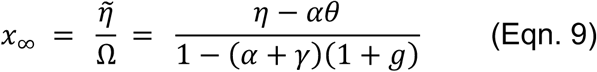

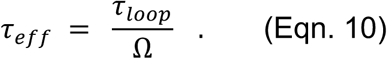

**Figure 3.**
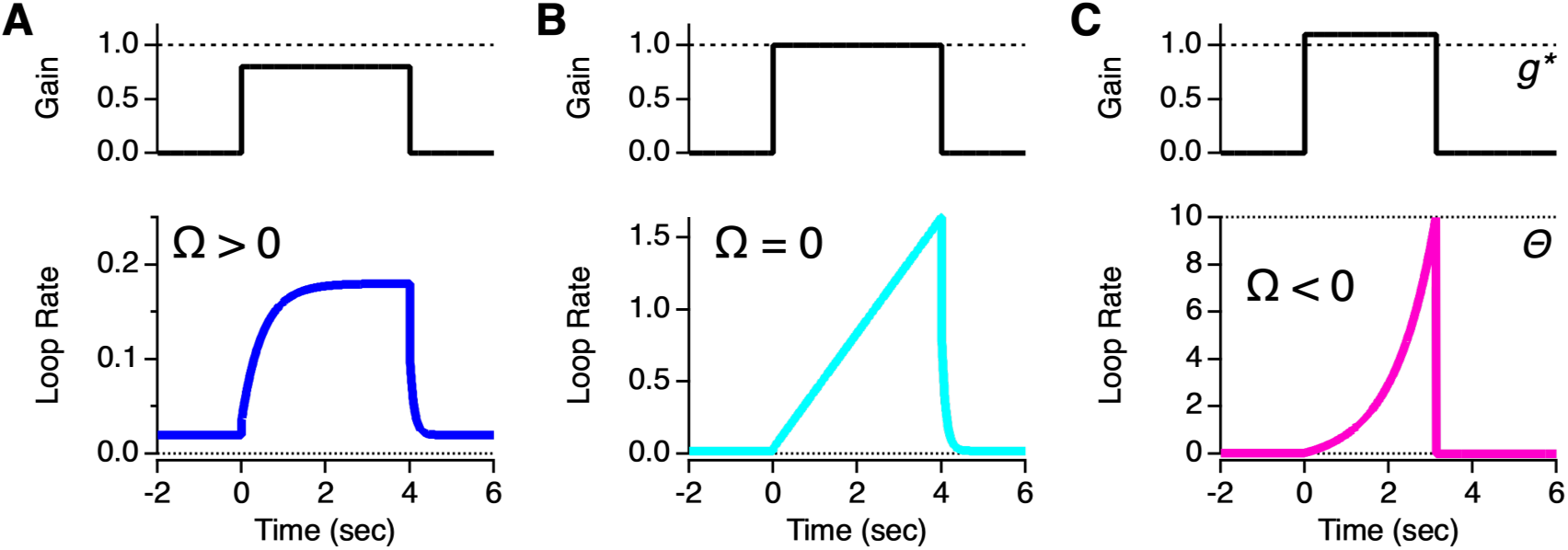
Different dynamical regimes of a single thalamocortical loop. **A.** For the case of Ω > 0, the firing rate in the layer 5 pool (blue; *below*) rises exponentially to a new steady-state level when descending cortical input (black; *above*) is turned on. **B.** For the case of Ω = 0, neural activity ramps linearly in time (cyan). **C.** For the case of Ω < 0, neural activity rises exponentially (pink) until the threshold for triggering movement is reached (dotted line). Parameters are: α = 0.1, γ = 0.4, η = 0.01, τ_pool_ = 50 ms; gains are [0.8, 1.0, 1.1] for A–C, respectively.

In the case of Ω < 0, activity increases exponentially in time with the same time constant (but with –Ω; see Figure 3C). This unstable limit is not very useful for evidence accumulation. Instead, one can think of this limit as a situation in which a decision to trigger the movement encoded by the L5 PT population in this TC loop is inevitable. Here, the circuit benefits from ramping to threshold very quickly – so that the movement can be executed as quickly as possible – and the exponential amplification aids in these dynamics.

The most interesting case is Ω = 0, where the circuit is a perfect neural integrator (Figure 3B). Here, the activity will ramp up linearly in time. This behavior mirrors the ramping seen in the readiness potential as well as in single neurons during decision-making. The rate of linear ramping (with no sensory input) is 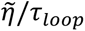.

Another interesting case is where the spontaneous input,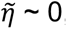, and the dominant input to the circuit is instead a sensory input, *S*(*t*). In this case, the circuit carries out an integral of the sensory input:

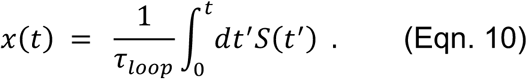

Because the case of the thalamocortical loop acting like a neural integrator is so interesting and potentially relevant, we consider what overall conditions make this possible. First, the loop should not have exponentially rising activity in the absence of cortical input. Such behavior would generate uncontrolled activity and would not be very useful. Thus, we require that α + γ < 1. This condition is in line with Crick and Koch’s hypothesis that there be no “strong loops” between thalamus and cortex (Crick & Koch, 1998).

Next, we would like the cortical input to layer 1, which raises the gain of the loop, to increase the loop’s feedback to achieve integrator dynamics. We also call this condition “ignition”, because activity in the loop will now be self-sustaining and increase in time due to its own dynamics. From this point-of-view, we can ask what amount of gain increase is necessary to trigger ignition, finding:

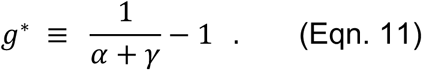

For greater concreteness, we choose values of the parameters of this model that are reasonable given what is known about the biophysics of this circuit. Experiments show that stimulation of the apical tuft of L5 PT pyramidal cells can turn a single somatic spike into a three spike burst (Larkum et al., 1999). Thus, the descending cortical command signal, *g*, is assumed to be roughly +100%.

A few notes about the parameter values. First, all firing rates are in “arbitrary units” that don’t have direct physical meaning. This is why it is natural to set the intrinsic gain with which voltage is converted into a firing rate equal to unity (*k* = 1). The time scale is set, in part, by the time constant of the cortical pool, τ*_pool_*. This has real time units (50 msec), so that we can take the time scales seriously. Also, 50 msec is roughly the period of the beta oscillation, which is prominent in the cortex during motor planning (Khanna & Carmena, 2015; Murthy & Fetz, 1996).

We choose the gain of the TC loop, α, to be 0.1. This means that the thalamus adds 10% of its firing rate to the cortical pool in every time step. While this value is not terribly well constrained by the literature, the current choice results in feedback that can have a significant effect without dominating the cortex. Similarly, the cross-inhibition is 0.1. This means that other TC loops subtract 10% of their firing rate from the given TC loop. This value is reasonable because lateral inhibition within the thalamus is also not completely winner-take-all. The recurrent feedback value of γ = 0.4 is gives rise to dynamics in the layer 5 pool with a time scale of 2τ*_pool_* = 100 ms.

Finally, the model has two more parameters. The spontaneous input is η = 0.01. We choose a small value, because the cortex exhibits a rough balance between excitation and inhibition (Dehghani et al., 2016; Shu et al., 2003). The movement threshold is Θ = 10. This high value allows us to better see the qualitative dynamics of the thalamocortical loop. Notice that here we are assuming that once the movement is initiated, the “goal” of higher cortical areas has been achieved, such that the gain input returns to zero. As a result, activity in the TC loop decays rapidly to its baseline value.

Overall, the individual thalamocortical loop serves as a variable gain amplifier. In the regime below ignition (Ω > 0), the TC loop responds in a graded fashion to sustained inputs. This could function as a form of “preparatory” activity that gives the loop a head start in competition with other loops (see below). In the regime above ignition (Ω < 0), the dynamics of the TC loop will rapidly ramp to threshold – potentially doing so faster than would be achieved without feedback from the thalamus. Thus, the extra excitation provided by the TC loop could be beneficial to help reduce the latency of movement. Finally in the integrator regime (Ω ∼ 0), the TC loop can integrate evidence over long timescales that are relevant to decision-making (see below). What is most important is that the same circuit can be rapidly transitioned from one regime to another under the control of descending cortical inputs, making it a highly flexible computational element.

***2) Two competing loops.*** Here, we again assume that activity in the thalamus is high enough to avoid truncation:

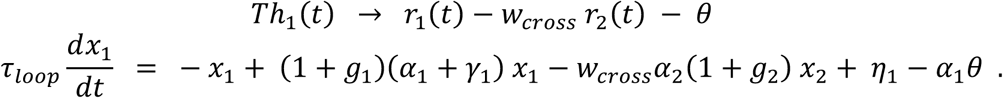

It is instructive to consider the (unlikely) case, where the two loops have identical properties and initial activity, α_1_ = α_2_, *g*_1_ = *g*_2_, *x*_1_ = *x*_2_. In this case, we get:

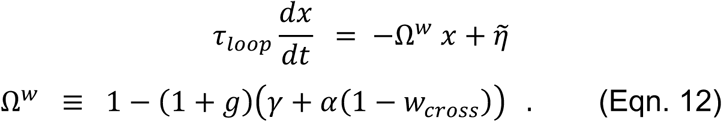

This has the effect of shifting the loop’s dynamics away from ignition. We can also define the effective decay for a single loop, Ω^0^, to explicitly see that cross-inhibition in the thalamus systematically increases the effective decay:

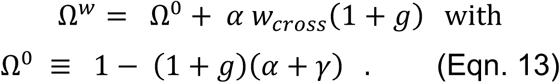

Alternatively, we can solve for the elevated gain input that is required to reach ignition:

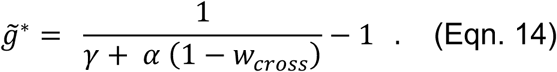

Generalizing to the case of *N* different loops all with such balanced activity, there is additional suppression given by *w*_*cross*_ → (*N* − 1)*w*_*cross*_. This result creates a mechanism in which activating too many TC loops makes it increasingly difficult for any of the loops to reach ignition and trigger a decision. We suggest in Discussion that this property may be related to the kind of indecision induced by having too many choices under consideration.

Another interesting limit is one in which both loops are individually close to the threshold of ignition, Ω_𝑙𝑜𝑜𝑝1_ ≈ Ω_𝑙𝑜𝑜𝑝2_ ≈ 0, but where the modified feedback is below threshold, 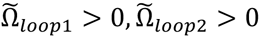. Here, we also assume that one of the loops has slightly stronger amplification, Ω_𝑙𝑜𝑜𝑝1_ > Ω_𝑙𝑜𝑜𝑝2_. This condition describes the case where two loops have nearly equal activity, but where one of the loops is poised to eventually win the competition.

In this case, the activity of both loops rises together (Figure 4A). This rise is slightly sublinear, because when both loops are equally active, neither loop is above the threshold for ignition. But because loop #1 has a slightly higher gain, this loop’s activity begins to exceed that of the other loop (time ∼ 1.5 sec). The difference increases over time, until loop #2 is driven down to a minimal firing rate. Once this happens, loop #1 reaches ignition and its activity ramps more steeply. (*Important note:* the timescale of this ramping activity is entirely dependent on the choice of model parameters; for instance, the models in figure 6 have shorter times to movement, chosen to match experimental data.)

**Figure 4.**
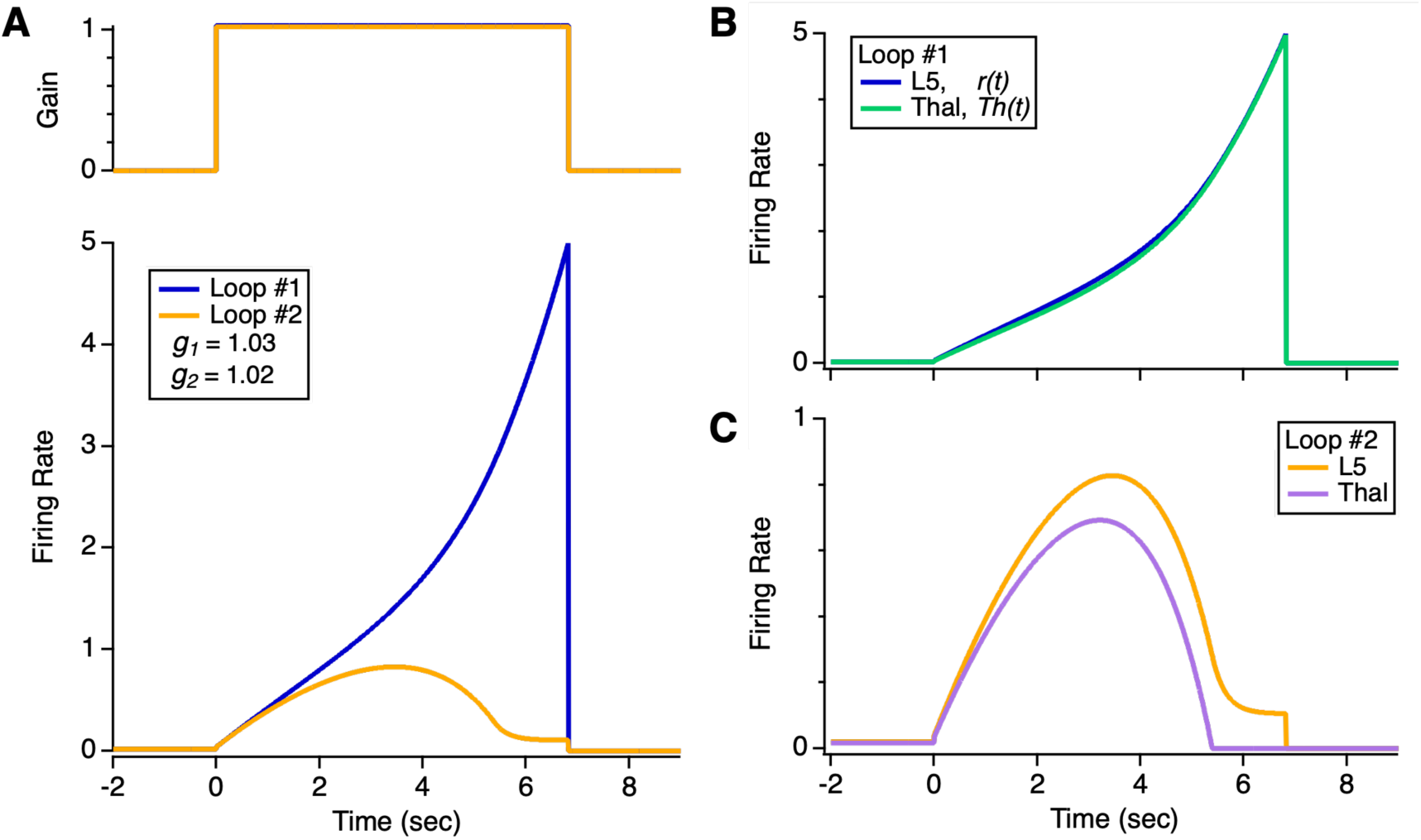
Two competing thalamocortical loops. **A.** *Top*: Cortical command signals for both loops versus time (blue vs. orange). Loop #1 has a slightly larger gain (*g_1_* = 1.03, *g_2_* = 1.02). *Bottom*: Firing rate of both loops versus time. Other parameters are [α = 0.1, η = 0.01, γ = 0.4, Θ = 5]. **B.** Firing rate of L5 cortical population (blue) and thalamus (green) for loop #1. **C.** Firing rate of L5 cortical population (orange) and thalamus (purple) for loop #2.

We can gain more insight into this process by comparing the firing rate of the L5 pool of cortical neurons versus that of the thalamic relay cells. For loop #1 (Fig. 4B), the thalamic firing rate is suppressed by a small amount early in the competition. But by the end, the difference vanishes. For loop #2 (Fig. 4C), the thalamic firing rate is more suppressed. Later in the competition, this firing rate is driven all the way down to zero. The cortical firing rate does not drop to zero, because of spontaneous inputs within the cortex.

### Which loop wins the competition?

As a general principle, the thalamocortical loop with higher activity will win the competition. This is because the activity in one loop inhibits activity in the other loop, which in turns disinhibits the first loop. Furthermore, if both loops are above the threshold for ignition, then their activity tends to increase over time. Thus, a small initial difference in activity between the loops will be amplified by winner-take-all dynamics into an increasingly large difference, until the loop with the higher activity reaches the threshold for movement. In this sense, the thalamus converts information from throughout the brain into a common currency for decision-making: namely, the firing of relay cells.

More specifically, we can ask how the parameters, [α, γ, *g*, η], determine which TC loop will win the competition. If the two loops have equal parameters, except for one, then the loop with the larger unique parameter will win. For instance, if loop #1 has stronger thalamic feedback, α_1_ > α_2_, with the other parameters being equal, then loop #1 will win. Similarly, for γ_1_ > γ_2_, *g*_1_ > *g*_2_, or η_1_ > η_2_. As these parameters combine to create an effective decay, Ω, one might think that the loop with the strongest amplification, *i.e.* the smallest decay, would win the competition. However, the amplification is changed by inhibition within the thalamus. As a result, amplification due to cortical gain control can be more effective than amplification due to thalamic feedback. In addition, the absolute firing rate also matters, because a loop that achieves higher activity can potentially suppress a loop with higher amplification. Overall, there is a complex and delicate balance between higher activity and higher amplification determining which loop wins the competition.

To illustrate this point, we will show several specific examples. In all cases, the TC loops are above the threshold for ignition, i.e. Ω_1_ < 0 and Ω_2_ < 0, so that their activity is self-sustaining, but below ignition in the presence of equal activity from the other loop, i.e. 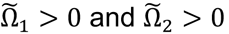.

In the first example, loop #1 has stronger thalamic feedback, but loop #2 has higher gain, such that both loops have the same effective decay, Ω_1_ = Ω_2_ = −0.2. Although both loops have the same intrinsic amplification, mutual inhibition in the thalamus reduces thalamic feedback from α to ∼α(1 − *w*_*cross*_). As a result, loop #1 has lower amplification in the presence of competition, and loop #2 wins (Fig. 5A). This example illustrates how a boost in amplification via cortical gain is generally more effective than a boost in amplification via thalamic feedback.

**Figure 5.**
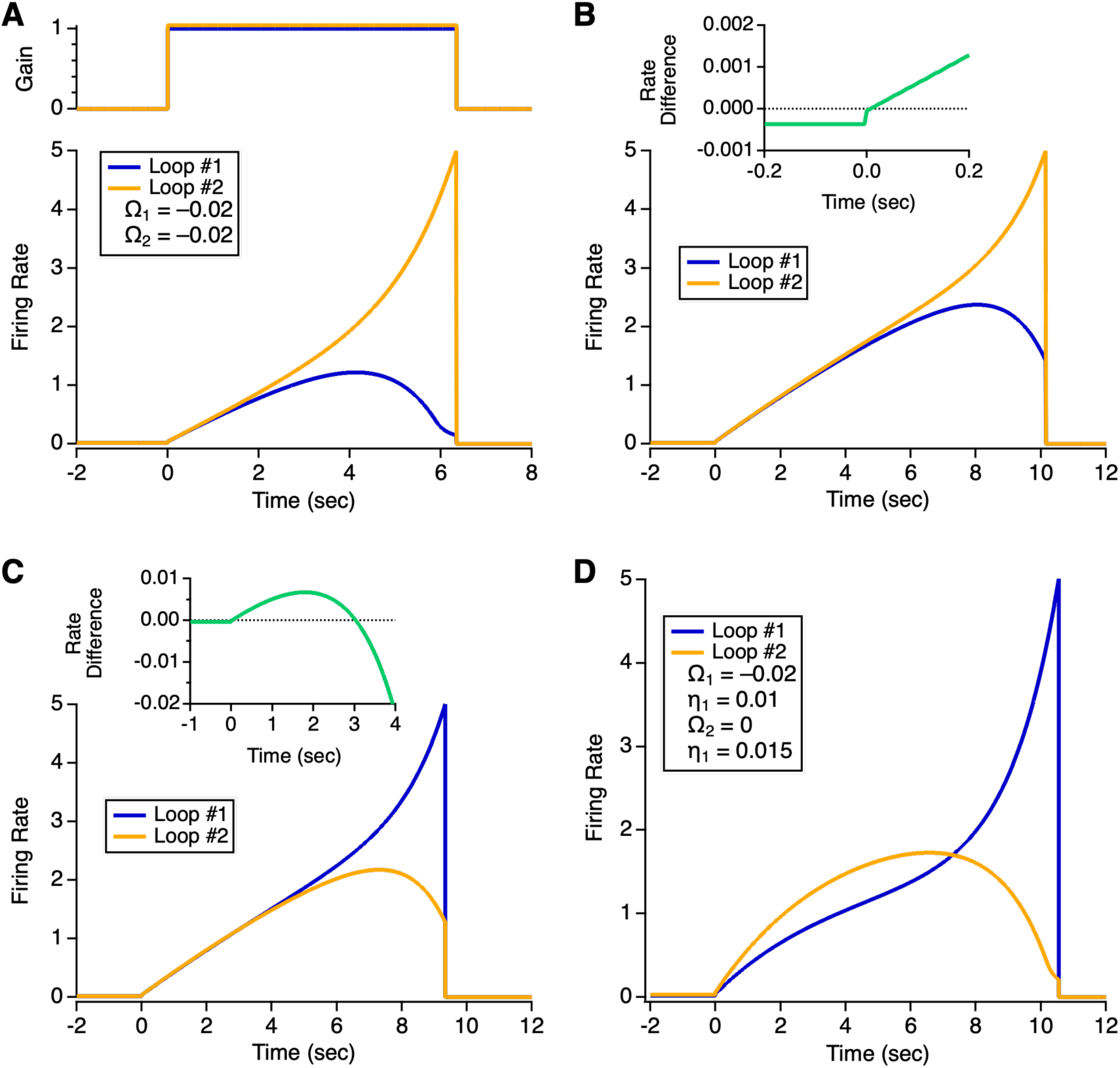
Factors that influence which loop wins the competition. **A.** *Top:* Cortical gain signals for two TC loops (blue vs. orange); *bottom*: firing rate for two TC loops; parameters are [α_1_ = 0.11, g_1_ = 1, α_2_ = 0.1, g_2_ = 1.04, η = 0.01, γ = 0.4]. **B.** *Top:* Firing rate difference (loop #2 minus loop #1) as a function of time relative to the onset of the cortical gain; *bottom*: firing rate for two TC loops (blue vs. orange); parameters as the same B, but with g_2_ = 1.0355. **C.** Same as B, but with different gain for loop #2, g_2_ = 1.035. **D.** Firing rate of two TC loops as a function of time after cortical gain turns on (blue vs. orange); parameters are [α_1_ = 0.11, η_1_ = 0.01, α_2_ = 0.1, η_2_ = 0.015, g = 1, γ = 0.4].

In this first example, we find that the amplification in the presence of competition favors 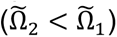. However, this criterion alone does not determine the winner. In the second example, loop #2 has a reduced gain, such that 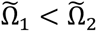. Despite loop #1 having greater amplification, loop #2 still wins the competition (Fig. 5B). We can gain some insight by zooming in on the difference in firing rate between the loops early in the competition (Fig. 5B *inset*). Before cortical gain is turned on, loop #1 has stronger thalamic feedback, so it starts with a higher steady-state firing rate. However, loop #2 has higher cortical gain, so once this gain turns on, loop #2 quickly achieves higher firing rate and ultimately wins the competition.

In the third example, the gain of loop #2 is reduced further. In this case, the activity in loop #2 is again larger after cortical gain is turned on, as in example #2. However, in this case, the amplification of loop #1 is sufficiently higher that its activity catches up to loop #2, eventually winning the competition (Fig. 5C). Together, these three examples illustrate the complex trade-off between higher activity and higher gain for competing thalamocortical loops.

For completeness, we show a fourth example. Here, loop #1 has stronger thalamic feedback, α_1_ > α_2_, and both loops have the same cortical gain. Instead, loop #2 has a higher spontaneous firing rate, η_2_ > η_1_. In this case, loop #2 has a higher steady-state firing rate before cortical gain is turned on. Furthermore, the activity of loop #2 greatly exceeds loop #1 in an early period after cortical gain is turned on. However, the greater amplification of loop #1 causes its activity to eventually catch up and ultimately win the competition (Fig. 5D). For slightly higher spontaneous firing rate, loop #2 instead wins the competition (η_2_ = 0.0152; *not shown*). These comparisons serve as additional examples of the complex trade-off between activity and amplification.

***3) Integration of sensory evidence.*** So far, we have assumed that the local input to layer 5 PT neurons is a constant, η. More generally, these neurons have apical dendrites that sample many synaptic inputs from layer 2/3, which exhibit extensive coding of sensory information. Thus, synapses made from layer 2/3 neurons onto layer 5 PT neurons can provide a time-varying sensory input, η = *S*(*t*). Then, when the thalamocortical loop is activated by descending cortical input, such that Ω ∼ 0, the loop can serve to integrate that time-varying stimulus. Furthermore, when the integrated activity reaches the threshold to activate the striatum, the movement encoded by that TC loop will be initiated. Thus, the thalamocortical loop can serve to integrate or accumulate sensory evidence and then the same TC loop can use this evidence to make the decision to move.

To explore this property, we consider an influential example of decision-making: the correlated dots motion task (Newsome & Pare, 1988). In this task, a monkey views a visual stimulus that consists of a fraction of dots that appear and move in the same direction together with a background of dots that appear and move in random directions. Coherence is defined as the fraction of all dots moving in the same direction. The task for the monkey is to make an eye movement that signals whether the coherent dot motion is to the right or left. Many neurons in cortical area MT are tuned for direction of motion and have a sustained firing rate during the random dot motion that is roughly proportional to the motion coherence (Britten et al., 1993). Neurons in hierarchically higher cortical areas, like area LIP, embody a temporal integral of the motion coherence (Roitman & Shadlen, 2002). These factors have led to the hypothesis that the brain computes a temporal integral over the activity of direction-tuned neurons in MT (Gold & Shadlen, 2007). While the neural mechanisms underlying this particular example of integration are not yet known, signatures of temporal integration have been seen in many experiments measuring neural activity and/or behavior (Brody & Hanks, 2016). Thus, the integration of sensory evidence appears to be a wide-spread computation in the brain.

For purposes of simulating evidence accumulation in TC loops, we abstract the details of these experiments and sidestep the controversies (Katz et al., 2016) (Latimer et al., 2015). We consider the sensory stimulus to be sampled from a Gaussian distribution with a mean, μ, a standard deviation, σ, and a correlation time, τ. Left-ward motion has a positive value, and rightward motion has a negative value. The coherence is defined as the ratio of the mean to the standard deviation of the motion stimulus, *C*≡ μ / σ. The stimulus is analyzed into leftward and rightward motion by direction-selective sensory neurons that compute instantaneous rectified-linear functions of the stimulus (Fig. 6A). The leftward-selective neurons form the sensory input to TC loop #1, *S_1_*(*t*), and the rightward neurons form the sensory input to TC loop #2, *S_2_*(*t*) (Fig. 6B).

At low coherence, activity in the two competing TC loops rises together (Fig. 6C), while at higher coherence, activity in the preferred loop rapidly suppresses the other (Figs. 6D). Furthermore, at low coherence, movement is triggered after a longer duration of stimulation than at higher coherence (Figs. 6C,D). The task performance – measured as the probability of selecting the correct motion direction – increases monotonically with a saturating function of coherence (Fig. 6E), and the movement time decreases monotonically with coherence (Fig. 6F), as seen in experiments (Roitman & Shadlen, 2002). Thus, the model of competing thalamocortical loops captures the primary qualitative features of this experiment involving evidence accumulation.

In order to explore the relationship between our model of competing thalamocortical loops and models of evidence accumulation, we formulated an explicit version of the drift diffusion model (DDM) (Mazurek et al., 2003; Ratcliff et al., 2016). Here, leftward and rightward motion (Fig. 7A *left*) are both passed through a perfect neural integrator that computes the integrated motion stimulus, scaled appropriately to compare against activity in the TC loops (Fig. 7A *middle*):

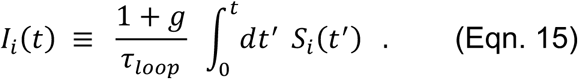

**Figure 6.**
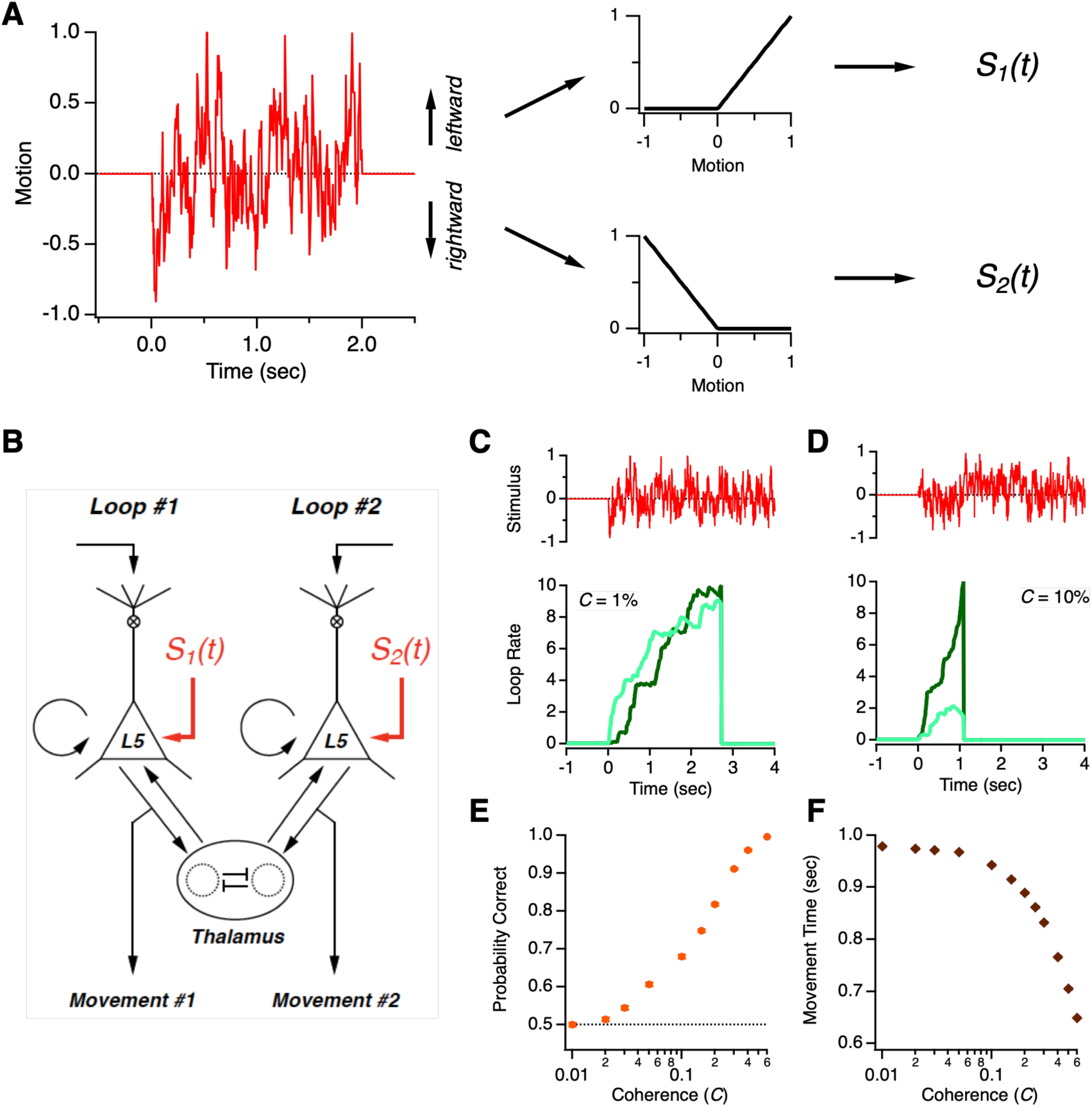
Integration of Sensory Evidence. **A.** A stochastic motion signal (*left*) is decomposed into neural channels representing leftward and rightward motion (denoted as *S_1_(t)* and *S_2_(t)*, *right*) by rectified linear transforms of the stimulus (*middle*). **B.** Two competing thalamocortical loops receive input from the two motion stimuli, *S_1_(t)* and *S_2_(t)*, respectively. **C, D.** *Above*: motion stimuli; *below*: activity of two competing TC loops (colors) for motion coherence = 1% (**C**) and 10% (**D**). **E.** Probability of correct decision plotted versus coherence. **F.** Time of movement decision plotted versus coherence. TC loops parameters are [η = 0.01, *g* = 1.03, *w_cross_* = 0.1, α = 0.1, γ = 0.4]. The stimulus has τ = 50 ms.

**Figure 7.**
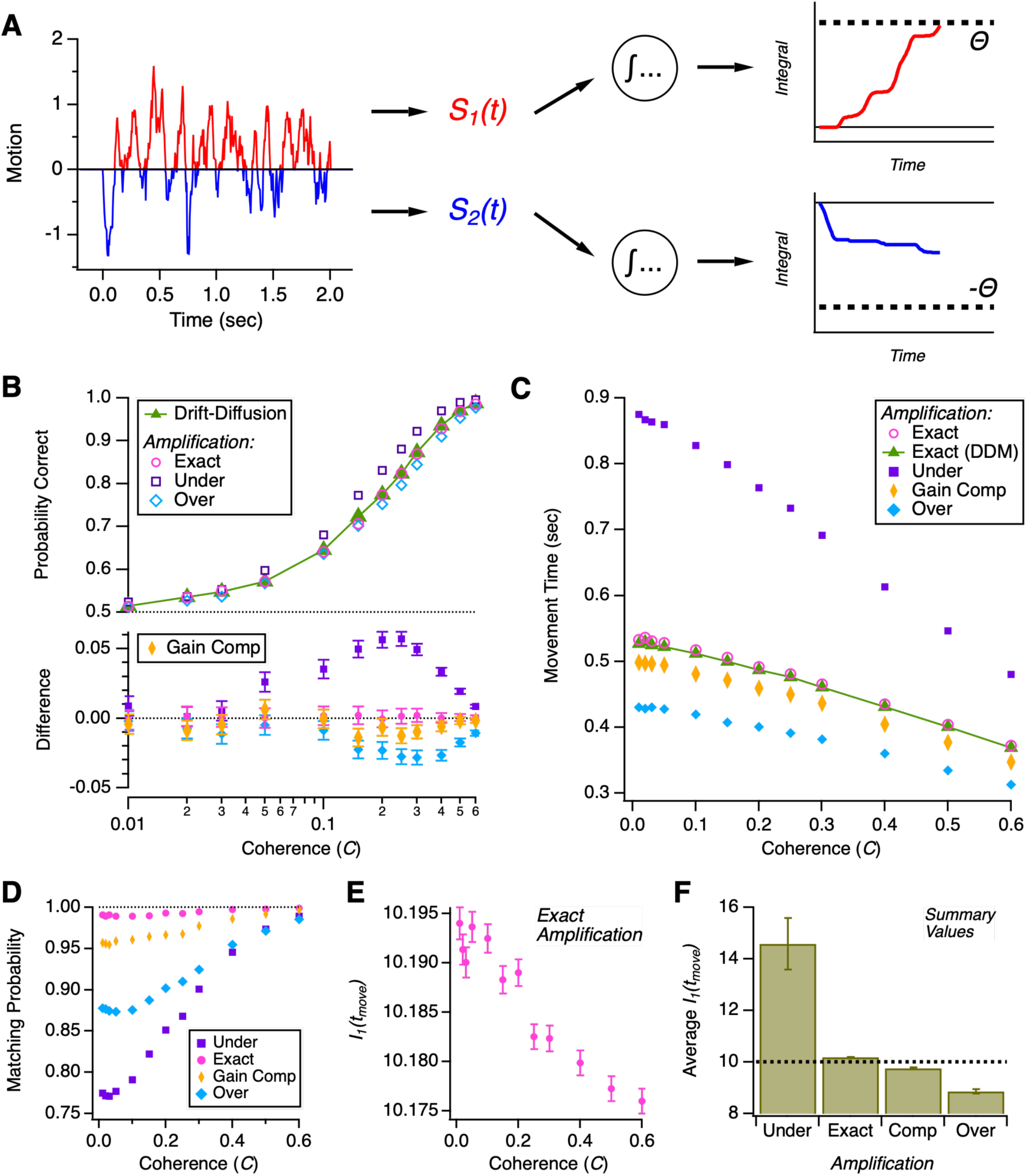
Comparison with the Drift Diffusion Model. **A.** In the drift diffusion model (DDM), leftward and rightward motion stimuli (*left*, colors) are separately integrated (*middle*) and compared against the same threshold (*right*). **B.** *Above*: Performance versus coherence for the DDM (green) compared against competing TC loops with three different sets of parameters (colors)*. Below:* Difference in performance between TC loops and DDM plotted versus coherence (colors). **C.** Movement time versus coherence for the DDM (green) compared against competing TC loops with different sets of parameters (colors). **D.** Probability of TC loops matching DDM decisions for different parameters (colors) plotted versus coherence. **E.** Integrated motion stimulus at the time of movement, *I_1_*(*t_move_*), plotted versus coherence. F. Summary of average value of *I_1_*(*t_move_*) for TC loops with different parameters. TC loop parameters for exact amplification: [η = 0, *g* = 1, *w_cross_* = 0.01], gain compensated: [η = 0, *g* = 1.0482, *w_cross_* = 0.1], under amplified: [η = 0, *g* = 0.9, *w_cross_* = 0.1]; over amplified: [η = 0, *g* = 1.1, *w_cross_* = 0.1].

Each integrated stimulus is then separately compared against the threshold (Fig. 7A *right*). Whichever integrated motion value reaches threshold first wins the competition and generates its corresponding movement. We first compare the DDM against a version of the competing TC loops model that comprises “exact” amplification. This model has a cortical gain input perfectly tuned to integrate its input, *Ω* = 0, and has no addition bias in its input, η = 0. Such a TC loop would, by itself, agree perfectly with the DDM. However, the competition between two loops via the thalamus introduces a deviation from the DDM. For exact amplification, we use a low value of the cross-inhibition in the thalamus, *w_cross_* = 0.01 (keeping in mind that prefect agreement only holds for *w_cross_* = 0). We found that the performance of the exact case agreed almost perfectly with the DDM (Fig. 7B). The time to movement was slightly longer (average value = 5.0 ± 1.0 ms), due to the non-zero cross-inhibition in the thalamus (Fig. 7C).

Next, we formulated a different version of the TC loops model. Here, we restore a more reasonable value of the cross-inhibition, *w_cross_* = 0.1, and note that early in the competition, when activity in both loops is closely matched, the effective decay is shifted from Ω to 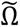. We compensate for this decreased amplification by boosting the cortical gain to 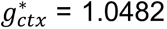, the value needed to reach ignition for effective gain 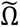 (Eqn. 20). The case of gain compensated amplification resulted in identical performance to the DMM (Fig. 7B) along with a slightly shorter time to movement (average = 17 ± 2 ms; Fig. 7C).

To better explore the sensitivity of parameters to sensory integration, we tested two other cases: under amplified (*g* = 0.9) and over amplified (*g* = 1.1). The underamplified TC loop actually exhibited slightly better performance than the DDM. This was accompanied by far longer times to movement. Presumably, the TC loops can achieve higher signal-to-noise ratio in integrating the noisy sensory stimuli in this case and thereby achieve better performance than an exact integrator. The overamplified case was the opposite: slight worse performance along with shorter times to movement.

We can gain more insight into the performance of competing TC loops by comparing their decisions with the DDM on a trial-by-trial basis by calculating the probability of their decisions matching (Fig. 7D). TC loops with exact amplification had very close agreement with the DDM (average = 99.3 ± 0.4%). The matching probability progressively degraded for the cases of: gain compensated, over amplified, and under amplified. It is interesting to juxtapose these results with performance itself (Fig. 7B). In particular, the under amplified TC loops achieved better performance while having a relatively poor match to the DDM. For all cases, the matching probability increased toward unity as the coherence increased.

Finally, we also calculated the integrated leftward motion in the DDM at the time at which the TC loops initiated a movement, *I_1_*(*t_move_*). For exact amplification, this value was close to the threshold, Θ = 10, with a trend towards a closer match for higher coherence (Fig. 7E). For under amplification, the integrated motion at movement was substantially higher than the threshold, consistent with the longer time to movement. For gain compensated and over amplified TC loops, the integrated motion at movement was below threshold, consistent with shorter times to movement.

Overall, this detailed comparison shows that not only do competing TC loops agree qualitatively with the drift diffusion model of evidence accumulation, but they agree in the task performance on a quantitative level over a broad range of parameters. Thus, competing TC loops have the potential to constitute a robust mechanism for the integration of sensory evidence towards movement. The time to movement varied more widely with the parameters of the TC loops. However, this variation gives rise to an interesting form of flexibility: TC loops can achieve better estimation performance at the cost of longer decision-making times via weaker than exact amplification, *or* the loops can achieve essentially equal performance with shorter decision-making times via stronger amplification.

***4) The impact of cortical feedback into layer 5.*** So far, we have assumed that the descending cortical input to layer 1 only changes the gain of firing in layer 5 PT pyramidal cells. This assumption is not strictly correct. First, inputs to the apical tuft will also depolarize the soma of L5 PT neurons (Larkum et al., 2004). Second, descending cortical axons often send collaterals into other layers of the local cortical circuit (Felleman & Van Essen, 1991), which would result in direct depolarization of the soma of L5 PT cells. This depolarizing input will be proportional to the strength of the cortical input. This simply augments the spontaneous input to the TC loop:

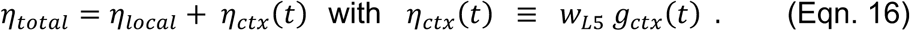

***5) The impact of thalamic feedback into layer 1.*** Axons from matrix-type thalamic relay cells that project to the neocortex synapse prominently in layer 1 (Clasca et al., 2012; Jones, 2001). The strength of this input will be proportional to the firing rate in the pool of thalamic relay cells but with a different proportionality constant than the input to layer 5. This input to layer 1 further boosts the gain of L5 neurons:

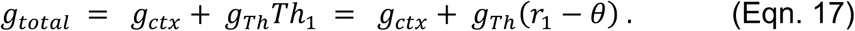

In principle, thalamic feedback to layer 1 will also have an “additive” effect on the firing of L5 neurons via direct depolarization. We can include this by adding an extra term to η*_total_*, which is ∼(𝑟_1_ − *θ*). However, this term has a mathematically identical effect as the original term for thalamic feedback, namely α (𝑟_1_ − *θ*). Thus, we can think of this effect as being absorbed into the parameter, α.

This boosted gain can also be characterized by having an effective decay rate, Ω^L1^, that systematically decreases as a function of the firing rate:

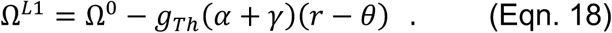

One way to characterize the effect of this extra feedback into layer 1 is to simulate the dynamics for increasing values of *g*_*Th*_. Because the input to layer 1 should generally scale with the strength of input to layer 5, we choose to parametrize this as *g*_*Th*_ = *f*α⁄*r*_0_. With these variables, the factor *f* is the ratio of thalamic contacts onto the apical tuft in layer 1 versus basal dendrites in layer 5, which is a property reported in the literature. In addition, we need another factor to describe the proportionality constant for how excitation from the thalamus changes the gain of L5 PT neurons. This proportionality has not been reported in the literature. For convenience, we choose *r*_0_ = Θ, so that the gain increases by one when the firing rate reaches the threshold to initiate movement.

Here, we show a case where the TC loop is below the ignition threshold with *g*_*Th*_ = 0 (Figure 8). For small values of *g*_*Th*_, the gain remains very close to its original value of *g_ctx_* = 0.9, and the firing rate saturates at a steady-state value (Fig. 8 *red*). As the feedback to layer 1 is increased, the gain begins to ramp up during the command period, and the firing rate is boosted to the point where it increases with roughly linear ramping (Fig. 8 *light blue*). At even higher feedback to layer 1, the ignition threshold is reached earlier in the command period, leading to a positive feedback loop between higher firing and higher gain. This produces a sharp ascent to the threshold for initiating movement (Fig. 8 *blue*). At even higher L1 feedback, this sharp positive feedback or “spike” occurs earlier (Fig. 8 *purple*).

**Figure 8.**
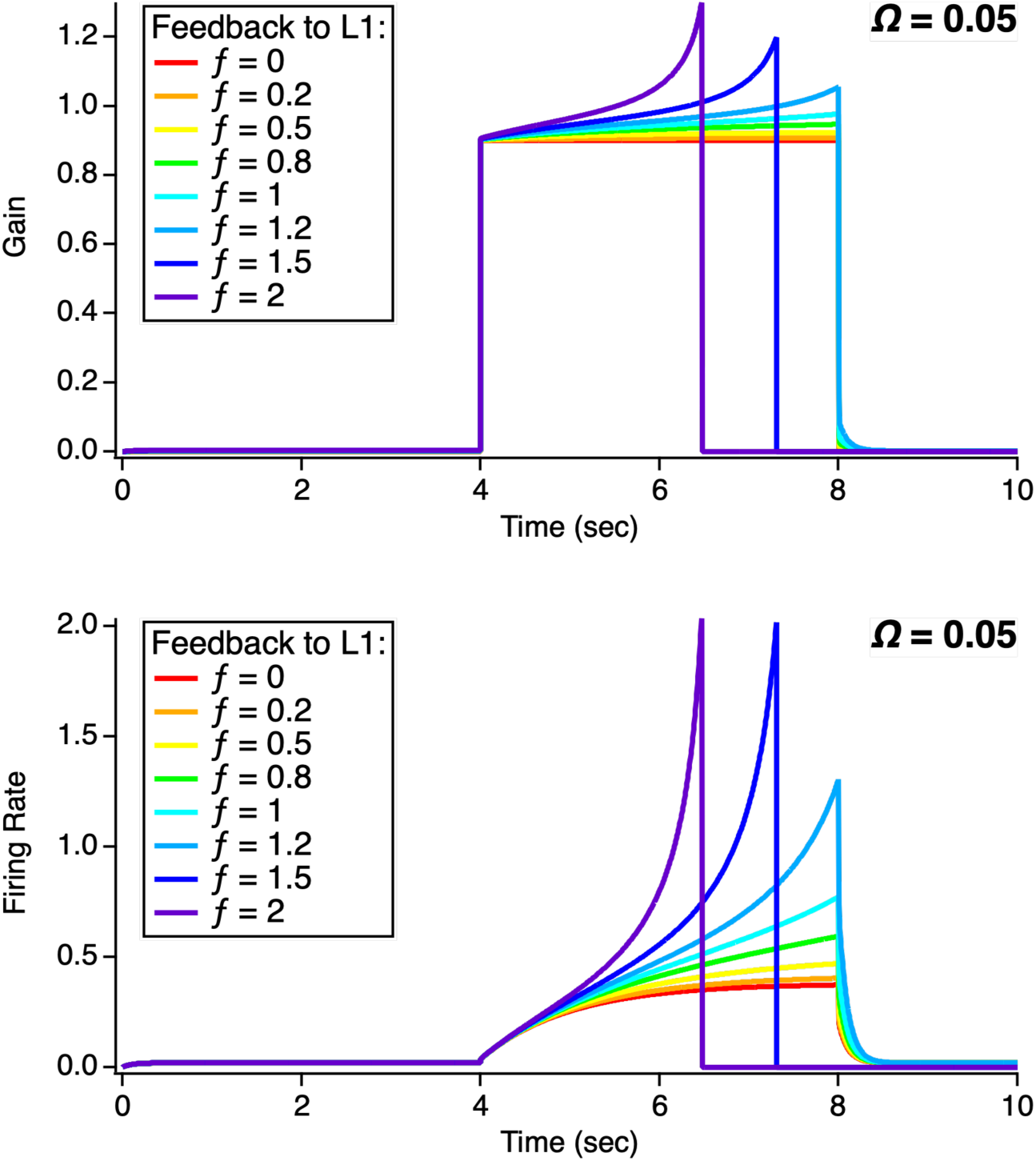
Thalamic feedback to layer 1. ***Top, Bottom***: (Gain, firing rate) of the thalamocortical loop as a function of time for increasing values of feedback from thalamus to layer 1, ƒ (colors), respectively.

This calculation illustrates that thalamic feedback into layer 1 boosts activity in the TC loop and effectively lowers the threshold for the cortical input required to reach ignition. But at the same time, strong feedback to layer 1 makes stable integration and accumulation of evidence difficult, as increasing activity in the TC loop can readily cause amplification in the circuit to rise above the level needed for integration. Therefore, we argue that from a functional perspective thalamocortical loops that participate in evidence accumulation should be in a limit where the feedback from thalamus to layer 1 has a small effect on the gain (*g_th_* ∼ 0).

On the other hand, the regime where thalamic feedback causes significant gain modulation of L5 PT neurons is interesting in a different way. As an example, we choose ƒ = 1 and vary the level of descending cortical gain control, *g_ctx_*. (Fig. 9). For particular values of the gain, the TC loop can remain poised at roughly constant gain and firing rate for long periods of time before sharply ramping to threshold. In all such cases, ramping to threshold emerges intrinsically from the dynamics of the thalamocortical loop without any additional external perturbation. Furthermore, the time required to reach the movement threshold diverges as the gain is tuned closer to the critical gain, 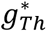 (Fig. 9C; see Eqn. 20 below).

**Figure 9.**
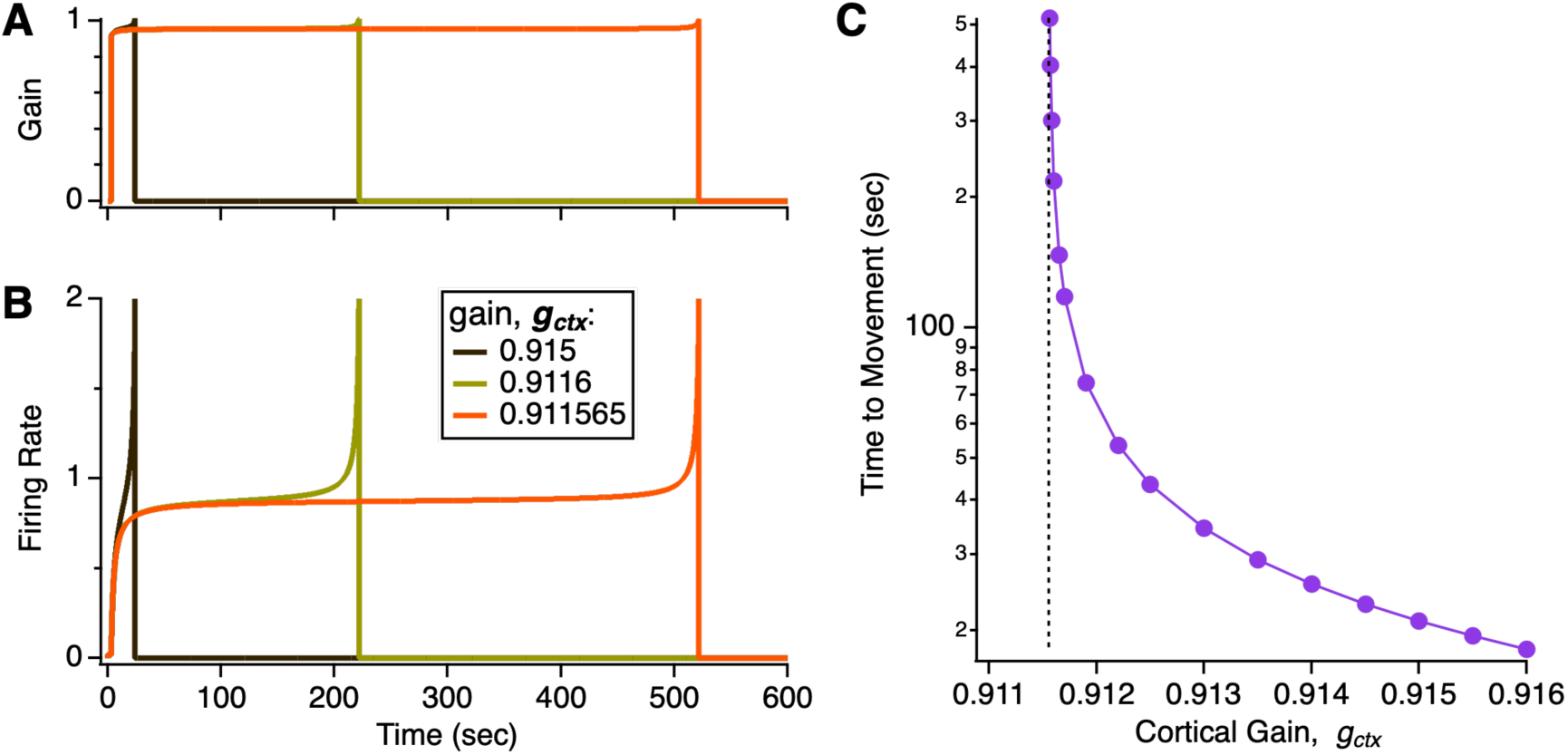
Thalamocortical loops can have long time scales to reach a decision. **A,B.** (Gain, firing rate) of a thalamocortical loop as a function of time for different levels of cortical feedback, *g_ctx_* (colors). **C.** Time until the thalamocortical loops reaches the threshold to trigger a movement plotted against cortical gain, *g_ctx_*; dashed line is the critical gain, 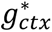 Other parameters are [α = 0.1, γ = 0.4, η = 0.01, Θ = 2].

We interpret this regime as one in which the TC loop can be thought of as making a “spontaneous decision”. This is because the firing rate of loop can appear to be stable for long periods of time and then rise sharply to threshold. Because of the delicate balancing required to remain in this stable regime, small excitatory perturbations will cause ramping to threshold to occur earlier. Thus, it is possible for the TC loop to have no change in its inputs, either in terms of cortical gain or sensory evidence, and yet for spontaneous fluctuations of firing within the cortical or thalamic neural populations themselves to cause a sudden rise to threshold. In this case, it would appear to the rest of the brain as if this thalamocortical loop had “decided by itself”. Therefore, we suggest that these dynamics in TC loops might contribute to the neural events that generate movements of an animal that are not triggered by external events (i.e., spontaneous movements).

Regardless of the strength of feedback to layer 1, we can derive some further analytic results that help understand the effects. We consider the case where the TC loop is below the threshold for ignition. In this case, the descending cortical input, *g_ctx_* > 0, causes the firing rate of the TC loop to rise to a new steady-state value. Using Eqn. 17 above for the total gain change, we can solve for the new steady-state voltage:

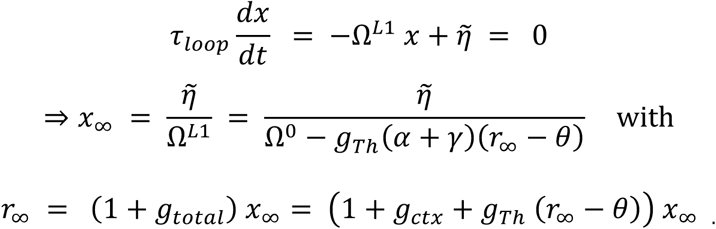

These self-consistent equations for the steady-state firing rate, *r*_-_, can be solved analytically. To simply its form, we assume θ = 0:

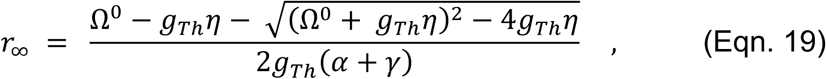

remembering that:

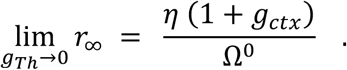

We can solve for how thalamic feedback into layer 1 boosts the steady-state firing rate (Fig. 10A). Notice that for weak L1 feedback, the effects are mild and scale roughly linearly. But for stronger L1 feedback, the increase in firing rate is super-linear. Finally, at a higher value, 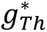, the feedback is strong enough to reach ignition, and thus there no longer is a solution to the steady-state firing rate, *r*_-_ (for example, in Fig. 10A *dark green*, 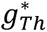, ≈ 0.064). As the cortical gain decreases, the steady-state firing rate decreases and larger values of *g*_*Th*_ are needed for ignition (Fig. 10A colors).

**Figure 10.**
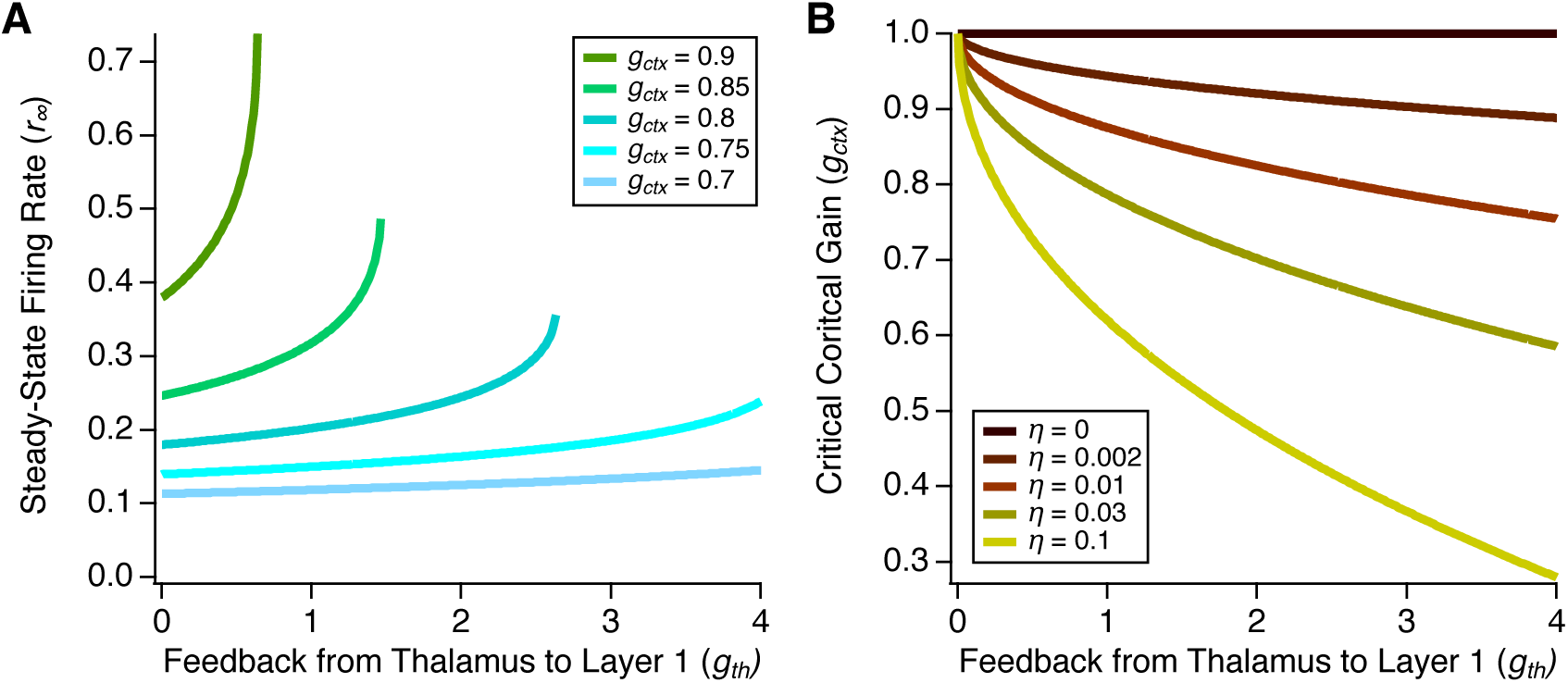
Analysis of thalamic feedback to layer 1. **A.** Steady-state firing rate, *r*_∞_, versus thalamic feedback to layer 1, *g*_*Th*_ for different values of cortical gain, *g_ctx_* (colors). **B.** Critical value of cortical gain input that leads to ignition, 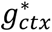 versus *g*_*Th*_ for different values of local input, η (colors). Other parameters are [α = 0.1, γ = 0.4, Θ = 1] and η = 0.01 (A).

Alternatively, we can ask how feedback into layer 1 changes the criterion for reaching ignition. In this case, ignition is achieved when there is no longer a real solution for the steady-state firing rate; this corresponds to the argument inside the square root in Eqn. 19 being negative. We can solve for the critical value of the cortical gain input that achieves ignition:

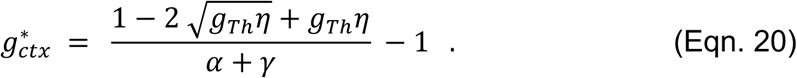

Here, we can see that the first term is the critical gain boost needed with no thalamic feedback to layer 1. Otherwise, stronger feedback to layer 1 decreases the cortical input needed to achieve ignition (Fig. 10B).

## Discussion

In summary, we have described a model of neocortical motor control, where each neocortical microcircuit can participate in the planning and generation of multiple motor outputs. Pools of layer 5 pyramidal tract (L5 PT) neurons that encode specific movements can exhibit motor planning dynamics, where their activity is enough to drive some subcortical targets, such as the thalamus, but not strong enough to drive the striatum sufficiently to suppress tonic inhibition in the pallidum and release their encoded movement. In this regime, matrix-type relay cells in higher-order thalamic nuclei feedback to the same L5 PT neurons, forming specific amplifier circuits. Activity in multiple thalamocortical (TC) loops interact via lateral inhibition in the thalamus, resulting in a winner-take-all dynamic among the competing TC loops. This winner-take-all dynamic then implements a form global selection among the motor plans of all the different neocortical microcircuits, allowing only a single motor plan to be expressed as movement and thus preventing inconsistent and dysfunctional activation of muscles and limbs.

The degree of amplification in thalamocortical loops is described by an effective decay constant, Ω, which is a function of the strength of thalamic feedback, α, the strength of local recurrent excitation, γ, and the strength of descending cortical feedback, *g*. We assume that the first two parameters are constant on a short time scale, such that the amplification of the TC loop is controlled dynamically by cortical feedback. Cortical feedback thus determines three regimes of TC loop dynamics: 1) a damped regime (Ω > 0), where activity in the TC loop reaches an elevated steady-state; 2) an overamplified regime (Ω < 0), where activity grows exponentially in time; 3) a balanced regime (Ω = 0), where neural activity is the time-integral of the synaptic inputs to the L5 PT neurons. Control by cortical feedback allows the TC loop to have stable, damped dynamics when its encoded movement is not involved in current motor plans but then rapidly switch into contributing to motor planning, as needed.

In the damped regime, activity in the TC loop can be elevated from baseline, which then biases that loop in favor of winning subsequent competitions with other loops. We interpret this as a form of anticipation or potentiation of a given movement, which cortical feedback can select. The overamplified regime produces activity that rapidly rises to the threshold of movement initiation. We interpret this regime as relevant to the situation where the brain has effectively decided on making the given movement and simply wants to implement that movement quickly. Most interesting is the balanced regime, where TC loops can integrate sensory information towards initiating their encoded movement. When multiple TC loops are in this balanced regime, their winner-take-all dynamic implements an accumulation of sensory evidence for decision-making, as has been studied widely. In this case, the integrator-like dynamics of TC loops implements evidence accumulation, and the threshold for movement initiation created in the striatum is the mechanism for making decisions among competing motor plans.

### Structure of the Model

A key element in our model is the fact that descending cortical input to layer 1 modifies the gain of L5 PT neurons. While this feature has been observed in neural data (Larkum et al., 2004; Larkum et al., 1999), relatively few circuit-level models include this form of gain control. Gain control is crucial for the ability to tune the dynamics of the TC loop among its different possible regimes. Our model captures this dependence through the fact that the effective decay, Ω, is an explicit function of the descending cortical gain, *g* (Eqn. 7). If instead, descending cortical input had only an additive effect on L5 PT neurons, then the amplification of the TC loop would remain the same, regardless of the strength of descending cortical input (see Eqn. 16). This circumstance would be undesirable for the brain, as a given TC loop would either never be excitable enough for self-sustaining activity or have sufficient excitability to be perpetually amplifying regardless of external inputs.

We showed that with the proper tuning of cortical gain, the TC loop has integrator dynamics (Ω ∼ 0). We found that this regime matched the drift-diffusion model of evidence accumulation closely over a range of cortical gain values (Fig. 6). This insensitivity to the cortical gain leads us to suggest that this method of creating a neural integrator circuit might be more robust to variations in synaptic weights than models based on a single recurrent neural network (Sebastian Seung, 1998). However, further studies are needed to make this comparison more precise.

This model has several qualitative features that are in broad agreement with neurophysiology. First, TC loops can exhibit ramping dynamics. This agrees with many experiments that have found ramping dynamics in both cortical neurons and thalamic relay cells (Catanese & Jaeger, 2021; Cisek & Kalaska, 2005; Li et al., 2015; Roitman & Shadlen, 2002; Tanaka, 2007). Second, if multiple TC loops ramp together, then their field potentials can sum to create a ramping potential on the scalp. This agrees with the observation of the readiness potential during spontaneous decisions to move (Libet et al., 1983; Schurger et al., 2021; Shibasaki & Hallett, 2006; Travers et al., 2021). Third, when multiple TC loops are in competition, their mutual inhibition within the thalamus progressively reduces the amplification of all of the loops (see Eqns. 12 and 13). With appropriate parameter values, this can prevent any of the loops from winning the competition, even when a single TC loop would have had enough amplification to reach threshold by itself. This case broadly resembles the observation that having too many choices can slow down decision-making and make it more “difficult” (Churchland et al., 2008; Haynes, 2009). Fourth, when the competition among TC loops resolves at a level of activity below the threshold to move, then the suppression of competing loops increases the amplification of the winning loop. This results in a steeper ramp to threshold (see Figure 3). This property resembles the steeper ramping of neurons observed just before movement in some experiments (Roitman & Shadlen, 2002) as well as in some measurements of the readiness potential (Haggard & Eimer, 1999; Maoz et al., 2019; Wen et al., 2018).

### Cortical Cell Types

In the body of the paper, we used a simplified description of cortical cell types by focusing on the role of L5 PT neurons. Layer 5 has a second prominent cell type, described variously as: i) “IT” because its axon travels via the corpus collusum to the other cerebral hemisphere (intratelecephalic); ii) “L5A” because it is found primarily in the upper half of layer; iii) “small” or “non-tufted” pyramids because they have little or no apical tuft in layer 1 (Molnar & Cheung, 2006; Shepherd, 2013). Importantly, L5 IT neurons have a major subcortical projection to the striatum. Because of this connectivity, L5 IT neurons will contribute to the strength of cortical input to the striatum that releases the inhibitory brake to allow movement. We have not explicitly modeled L5 IT neurons here; instead, we assume that there will be a subset of L5 IT neurons that encode for a specific movement and that the firing of these neurons will be strongly correlated with the firing of L5 PT neurons that encode the same movement. Therefore, when the firing rate variable, *r*(*t*), reaches the striatal threshold, Θ, this threshold value incorporates the corresponding input from L5 IT neurons.

L5 IT cells have several other important properties: i) they are far more likely to be presynaptic to L5 PT neurons than vice versa; ii) feedback from matrix-type thalamic relay neurons is made substantially to layer 5A (Clasca et al., 2012; Harris & Shepherd, 2015; Hooks et al., 2013); iii) they project both to ipsilateral and contralateral striatum, while L5 PT neurons project only to the ipsilateral striatum (Shepherd, 2013). Considering these properties together, it is clear that L5 IT and PT cells must be closely coordinated. First, L5 IT neurons provide much of the presynaptic input to L5 PT neurons, so that clusters of activity in PT neurons might just be a downstream consequence of clusters of IT neuron activity. Second, thalamic feedback significantly targets IT neurons, so that the effective feedback from thalamus to L5 PT neurons, α, includes disynaptic excitatory pathways via L5 IT neurons. Third, any movements that require coordination across cerebral hemispheres must obviously rely on L5 IT neurons to tie together TC loops in both hemispheres.

Another major input from the cortex to the thalamus comes from L6 CT neurons (Jones, 2001; Shepherd & Yamawaki, 2021). These neurons provide modulatory input to thalamic relay cells, which may make only a mild contribution to total cortical excitation (Sherman, 2016). However, L6 CT neurons have extensive collaterals in the thalamic reticular nucleus (TRN), whose neurons are GABAergic and project widely onto relay cells (Halassa & Acsady, 2016; Jones, 2002). As a result, this pathway will contribute to lateral inhibition in the thalamus. We again assume that L6 CT neurons will have clusters of activity that encode specific movements and that these clusters are correlated with the respective clusters of L5 PT neurons (Guo et al., 2020). Therefore, L6 CT neurons will add inhibition between TC loops, which is incorporated into the parameter, *w_cross_*.

Recent studies of cell types in layer 5 have revealed two distinct populations of L5 PT neurons in the mouse motor cortex (Economo et al., 2018; Tasic et al., 2018). One population projects primarily to the thalamus and is located in the upper portion of layer 5B (PT_upper_), and the other population projects strongly to the medulla and is located in lower portion of layer 5B (PT_lower_). PT_upper_ neurons were found to have greater activity and task selectivity during motor preparation, while PT_lower_ neurons had greater activity and selectivity related to movement itself (Economo et al., 2018). In this context, our model resembles more closely PT_upper_ neurons. Furthermore, when the TC loop involving PT_upper_ neurons reaches threshold, it might trigger the activity of a subsequent population of L5 PT_lower_ neurons, which themselves generate movement, instead of directly triggering movement. One reason that this organization might be beneficial stems from the fact that in our model the dynamics of the TC loop is one-dimensional. Thus, when it reaches threshold, it is well-suited to trigger a single, ballistic movement – such as a saccade or a lick – but not an entire sequence of movements – such as reaching with the arm and grasping with the hand. If the TC loop that integrates evidence and “decides” in favor of a movement instead passes control off to a different population of L5 PT neurons, then this population can have synapses structured to generate internal dynamics to drive an extended sequence of individual movements.

At the same time, a few caveats about this distinction are worth mentioning. First, both PT_upper_ and PT_lower_ neurons project to the superior colliculus, which is a major motor organizing center that is already high in the motor hierarchy (Swanson, 2000). Thus, PT_upper_ neurons still have the ability to directly influence a wide variety of movements beyond just saccades and licks (Benavidez et al., 2021; Liu et al., 2022). Second, there is an extensive literature claiming that L5 PT axons innervate both the thalamus and medulla (Harris & Shepherd, 2015; Prasad et al., 2020; Shepherd, 2013), including studies based on single axon reconstructions (Deschenes et al., 1994; Kita & Kita, 2012). In addition, a comprehensive genetic analysis of cell types in both V1 and ALM showed that the markers that distinguish PT_upper_ versus PT_lower_ are present in ALM but not V1 PT neurons (Tasic et al., 2018). Therefore, this distinction might be present only in some cortical areas.

### Thalamic Input to Cortical Layer 1

In general, matrix-type relay cells send their densest projections into layer 1 as well as layer 5 (Clasca et al., 2012; Hunnicutt et al., 2014). In fact, a specific measurement of EPSCs in different cortical layers generated by thalamic input showed that monosynaptic inputs to layer 1 were ∼2-4 times stronger than inputs to layer 5 (Guo et al., 2018). However, any additive inputs to the somas of L5 PT neurons have mathematically an identical effect regardless of the layer in which they arrive (see section 5). Therefore, our definition of the strength of thalamic feedback, α, includes additive inputs, as measured by (Guo et al., 2018), from all layers. The important mathematical difference is whether thalamic feedback has an additive or multiplicative effect on the firing of L5 PT neurons.

The model we presented here used a very simple assumption for how thalamocortical feedback changes the gain of L5 PT neurons – namely, that this gain increases linearly as a function of the firing rate of thalamic relay cells (Eqn. 17). However, the increase in gain of L5 PT neurons appears to be mediated by the generation of Ca^++^ spikes in their apical tufts (Larkum et al., 2004; Larkum et al., 1999). This suggests that there is a threshold of thalamic activity that is required to begin to change L5 gain. Furthermore, the gain change is also expected to saturate at a maximal gain when Ca^++^ spikes are being robustly generated. Thus, a more realistic model of how thalamic feedback affects L5 gain might be a sigmoidal function of thalamic firing rate, not linear.

This more realistic control of L5 gain would have some important consequences for thalamocortical dynamics. First, a threshold for changing L5 gain creates a regime of more stable integration of sensory evidence, in which the thalamic firing rate was below this threshold to change L5 gain. Second, the saturation of L5 gain creates the possibility of bistable dynamics in the TC loops. In this case, a brief input to the TC loop could raise the L5 gain to a level that results in an elevated firing rate of the TC loop, which is then dynamically stable because the L5 gain saturated at a value below the threshold for ignition. Furthermore, the specific manner in which inputs to the apical tuft change L5 gain depends on the action of Martinotti (SOM^+^) cells (Murayama et al., 2009) as well as H-currents in the apical tuft (Guo et al., 2018). This opens up the possibility that the regulation of L5 gain by thalamic feedback could itself by tuned by modifying the activity of Martinotti cells or by having different densities of these cells in different cortical areas as well as by neuromodulatory control of the H-current. The issue of how a more realistic control of L5 gain affects thalamocortical dynamics is a topic that requires further investigation.

### Thalamic Cell Types

The model we have formulated here is best matched to the multiareal matrix cell type, which is the primary relay cell type in the motor thalamus (Clasca et al., 2012). Other thalamic nuclei, including the mediodorsal (MD) and the pulvinar (Pulv), predominantly have the focal relay cell type (Clasca et al., 2012). The multiareal cell type has significant collaterals in layer 5 as well as branches connecting to multiple cortical areas. In contrast, the focal cell type has spatially localized collaterals in layer 3 that are typically found in one cortical area. In the context of our model, the focal cell type would have a much lower value of the thalamic feedback parameter, α. At the same time, the connections to layer 3 can serve to increase the gain of L5 PT neurons (Peron et al., 2020; Quiquempoix et al., 2018), implying that firing of these focal-type matrix cells would have a greater impact on the gain of L5 PT neurons than does the firing of multiareal-type thalamic matrix relay cells. This makes the focal-type relay cell potentially well suited for implementing “rules” in higher cortical areas, like the prefrontal cortex (PFC) – a topic that we will return to in a later publication.

Cortical layers 2 and 3 are generally believed to play a major role in processing sensory stimuli, given its position in the neocortical microcircuit (da Costa & Martin, 2010; Harris & Mrsic-Flogel, 2013). This means that input from focal-type thalamic matrix cells may also trigger specific sensory representations. While these thoughts are speculative, we can propose two potential roles for this pathway. First, the thalamic feedback could boost the activity of sensory representations in layer 2/3 that serve as the sensory cue for activating the very same TC loop. In this case, the circuit would have enhanced amplification in a manner that is stimulus specific. This function would require that learning rules had previously shaped synaptic connectivity to enhance the specific pathways from focal-type relay cells → L3 → L5 PT. Second, multiple such TC loops might be able to compete via lateral inhibition in the thalamus in a manner that implements a winner-take-all dynamic that is specific to the sensory stimuli corresponding to each loop. This dynamic might implement a form of attentional control, thus giving the thalamus a role in attentional modulation, as has been previously proposed (Halassa & Kastner, 2017).

### Other Theories of Thalamic Function

The thalamus has traditionally been thought of as the “sensory gateway” to the neocortex. This is quite clearly the role of first order thalamic nuclei with core-type thalamic relay cells, such as the lateral geniculate nucleus (LGN), medial geniculate nucleus (MGN), ventral posterolateral (VPl) and ventral posteromedial (VPm) nuclei. These nuclei all connect subcortical sources of sensory information (vision, audition, somatosensation, respectively) with primary sensory cortices (V1, A1, and S1, respectively). Sherman suggests that higher-order thalamic nuclei continue these sensory pathways up cortical hierarch (Sherman, 2016, 2017). For instance, V1 projects to the pulvinar, which then projects to V2, which projects to a distinct region of the pulvinar, which projects to V4, etc. These pathways may form a sensory channel that parallels the traditional neocortical hierarchy.

Here, we propose that many of these pathways may fundamentally have a role in motor planning. The core of our argument is that fact that the driver-type inputs to higher order thalamus come from a set of neocortical neurons that send collaterals to brainstem motor centers (Guillery & Sherman, 2002). In this context, it is important to note that many neurons in the pulvinar have no visual receptive field, and those that do tend to have larger, less-specific receptive fields than neurons in corresponding areas of the neocortex (Chalupa & Abramson, 1989; Purushothaman et al., 2012). Thus, it has been argued that higher-order thalamic nuclei, like the pulvinar, are unlikely to serve primarily as a sensory relay (Halassa & Kastner, 2017). In addition, even random inputs from L2/3 to L5 PT neurons are likely to create some kind of a sensory receptive field (Hansel & van Vreeswijk, 2012), consistent with the observation of orientation tuning in layer 5.

Halassa has argued that the thalamus should be thought of as an amplifier of local cortical activity that can enhance information represented in the cortex (Schmitt et al., 2017). This proposal emerged from experiments in which silencing MD drastically reduced activity in PFC as well as reduced the ability of mice to follow behavioral rules. Our model is broadly consistent with this function, because: 1) feedback from the thalamus contributes significantly to ongoing cortical activity, 2) this feedback enhances specific subsets of cortical neurons, in this case representing different motor plans. This idea of the thalamus amplifying cortical activity was subsequently expanded to a more general role in attentional control (Halassa & Kastner, 2017). As mentioned above, attentional control is a function that can potentially be captured when our framework is extended to describe the role of focal-type thalamic matrix relay cells.

Another proposal is that the thalamus can restructure the dynamics of recurrent circuits in the neocortex when specific subsets of matrix-type relay cells are freed from inhibition from the basal ganglia (Logiaco et al., 2021). The thalamus can have this action by effectively adding a set of effective weights between cortical neurons that reflex the pathway from cortex to thalamus and back. This is a substantially different proposal than our model. However, Logiaco et al. also propose that such motifs can be rapidly selected by activation of different “preparatory” thalamocortical loops. These preparatory loops share many qualitative features with our model of TC loops participating in motor planning via evidence accumulation. Furthermore, one could also imagine identifying preparatory loops with PT_upper_ neurons in layer 5 and the dynamics shifting circuits with PT_lower_ L5 neurons (Economo et al., 2018).

